# A low dimensional cognitive-network space in Alzheimer’s disease and frontotemporal dementia

**DOI:** 10.1101/2022.08.29.504748

**Authors:** Lorenzo Pini, Siemon de Lange, Francesca Pizzini, Ilaria Boscolo Galazzo, Rosa Manenti, Maria Cotelli, Samantha Galluzzi, Maria Sofia Cotelli, Maurizio Corbetta, Martijn Van den Heuvel, Michela Pievani

## Abstract

Network neuroscience is a promising approach to explore cognitive processes in neurological disorders. Alzheimer’s disease (AD) and frontotemporal dementia (FTD) show network dysfunctions linked with cognitive deficits. Within this framework, network abnormalities between AD and FTD show both convergent and divergent patterns. However, these functional patterns are far from being established and their relevance to cognitive processes remains to be elucidated. In this study, we aimed to investigate the relationship between cognition and functional connectivity of major cognitive networks in these diseases. Twenty-three bvFTD (age: 71±10), 22 AD (age: 72±6) and 20 controls (age: 72±6) underwent cognitive evaluation and resting-state functional MRI. Principal component analysis was used to describe cognitive variance across participants. Brain network connectivity was estimated with connectome analysis. Connectivity matrices were created assessing correlations between parcels within each functional network. The following cognitive networks were considered: default mode (DMN), dorsal attention (DAN), ventral attention (VAN) and frontoparietal (FPN) networks. The relationship between cognition and connectivity was assessed using a robust convergent correlation-wise and interaction analyses. Three principal cognitive components explained more than 80% of the cognitive variance: the first component (cogPC1) loaded on memory, the second component (cogPC2) loaded on emotion and language, the third component (cogPC3) loaded on the visuo-spatial and attentional domains. Compared to HC, AD and bvFTD showed impairment in all cogPCs (p<0.002), and bvFTD scored worse than AD in cogPC2 (p=0.031). At the network level, the DMN showed a robust association in the whole group with cogPC1 and cogPC2, and the VAN with cogPC2. By contrast, DAN and FPN showed a divergent pattern between diagnosis and connectivity for cogPC2. We confirmed these results by means of a multivariate analysis (canonical correlation). These results suggest that a low-dimensional representation can account for a large variance in cognitive scores in the continuum from normal to pathological aging. Moreover, cognitive components showed both convergent and divergent patterns with connectivity across AD and bvFTD. The convergent pattern was observed across the networks primarily involved in these diseases (i.e., the DMN and VAN), while a divergent FC-cognitive pattern was mainly observed between attention/executive networks and the language/emotion cognitive component, suggesting the co-existence of compensatory and detrimental mechanisms underlying these components.

## 1. Introduction

Neurodegenerative disorders are among the top leading cause of death and disabilities worldwide^1^. Among people aged more than 65 years old, Alzheimer’s disease (AD) is the most common form of neurodegeneration, while frontotemporal dementia (FTD) represents the first cause of cognitive impairment in younger individuals. In AD, memory is typically the earliest sign of cognitive deterioration. FTD serves as an umbrella term for several clinical syndromes, including the behavioral variant FTD (bvFTD), usually characterized by behavioral disturbances in the earliest stages^2,3^. To date, no cure is available for these diseases. Recent significant advancement in the pharmacological field has been done, although findings are still far from being conclusive^4–6^. A better understanding of the pathophysiological mechanisms underlying these cognitive/behavioral symptoms might pave the way to novel treatments and rehabilitation options^7^.

Resting-state functional magnetic resonance imaging (rsfMRI) is widely used to assess the putative functional architecture of the brain at rest. This technique can investigate *in vivo* brain oscillations in the blood oxygen level dependent (BOLD) signal between different brain regions. Brain areas showing temporal BOLD synchronization are assumed to be functionally grouped into neural networks. Functional connectivity (FC) exhibits a low-dimensional spatiotemporal pattern^8,9^. This functional scaffold might have a representational role for cognitive abilities^10,11^. The default mode network (DMN) is associated with episodic memory performance and shows a gradual shrinking with aging, in line with the natural decline of memory performance in the elderly^12^. Similarly, a group of “attentional networks” is linked with executive, language, and attentional abilities, that is the frontoparietal (FPN), the dorsal attention (DAN), and the ventral attention (VAN) networks.

In typical AD, breakdown of DMN is linked with core symptoms, i.e. impaired episodic memory^13^. By contrast, bvFTD manifests reduced FC of the VAN (also referred to as salience network) that is linked with clinical severity^14^. Other cognitive functions and networks are involved during disease progression, such as attentional networks/functions in both conditions^15,16^. However, a simple 1:1 relationship between (lower) FC and (impaired) cognition is too simplistic to explain the complex pattern of cognitive and brain changes. Large-scale networks are closely interconnected and alterations in one network can have effects on other networks and undermine the balance of this functional scaffold. The triple network theory states that aberrant dynamic cross-network interactions of the VAN, FPN, and DMN underlie a wide range of cognitive/behavioral disturbances^17^. This theory posits that VAN integrates external information acting as an interface between DMN and FPN, regulating their competing inter-network activity and promoting appropriate behavioral response. Similarly, the VAN acts as a circuit breaker when attention is reoriented to relevant environmental stimuli, interrupting ongoing activity in the DAN, which in turn shifts attention to the new source of information^18,19^. These studies suggested a general role in switching between networks supporting cognitive functions, which may explain previous evidence of between-network alterations in neurological disorders. Brain stroke lesions increase connectivity between networks commonly anti-correlated, such as the DMN and the DAN, with detrimental consequences on cognitive abilities^20^. In neurodegenerative disorders, the pivotal study of Zhou et al. (2010) reported a divergent connectivity pattern in AD and bvFTD, whereby reduced connectivity of the DMN in AD was accompanied by hyper-connectivity of the salience network, while the opposite was seen in bvFTD. More recently, the same group observed that AD and bvFTD show divergent abnormalities in the topological organization of functional brain networks extending into subcortical and inter-network connections^21^. These studies pointed out a complex pattern of network connectivity alterations within the connectivity gradient. However, the relationship between this divergent functional pattern in AD and bvFTD and cognition is still unclear. A “classical” cognitive-network approach, which considers the relationship between a single test score (or a composite score across *apriori* defined domains) with FC might mask some latent relationships, considering also that cognitive scores are highly correlated. This phenomenon has been widely assessed in stroke where a low dimensional space can explain a large variance of cognitive deficits and is coupled with FC^20,22,23^. Based on these premises, in the present study we sought to investigate for the first time the relationship between cognitive deficits and FC within a new low dimensional space in the continuum of aging and age-related diseases. To this aim we used (i) a principal component analysis (PCA) to identify low dimensional representations of cognition and (ii) a connectome analysis to assess FC patterns in core cognitive networks. We then assessed convergent and divergent cognitive-connectivity relationships using univariate and multivariate analyses. We hypothesize to find, within a low-dimensional space, both shared and divergent cognitive-FC patterns.

## 2. Methods

### 2.1 Participants and study design

Patients were enrolled at the IRCCS Istituto Centro San Giovanni di Dio Fatebenefratelli in Brescia (Italy) as part of the NetCogBS project^24^ (ClinicalTrials.gov identifier NCT03422250). Patients underwent a clinical, cognitive, and imaging assessment. Cognitive and imaging variables were collected also in a group of age-matched healthy controls (HC). The study was conducted in accordance with the Declaration of Helsinki principles and approved by the local ethics committee of the IRCCS Istituto Centro San Giovanni di Dio Fatebenefratelli in Brescia (Italy). Written informed consent was obtained from all participants.

We included patients with a clinical diagnosis of AD or bvFTD^3,25^. Inclusion criteria for patients were: i) age between 50 and 85 years; ii) ability to provide written informed consent; iii) availability of a collateral source. We excluded patients with moderate/severe dementia (Mini-Mental State Examination (MMSE) score < 18), any medical condition that could interfere with assessments, and contraindications for MRI (metal implants, pacemakers, prosthetic heart valves, claustrophobia). Inclusion criteria for HC were a normal neuropsychological performance on the cognitive battery, with no personal history of neurological, psychiatric, or cerebrovascular disorders. Exclusion criteria are reported in Pini et al.^24^. For the present study, we included all the patients with available cognitive assessment and MRI examination performed at the baseline. The whole procedure of analysis is depicted in **Figure 1**.

**Figure 1.**
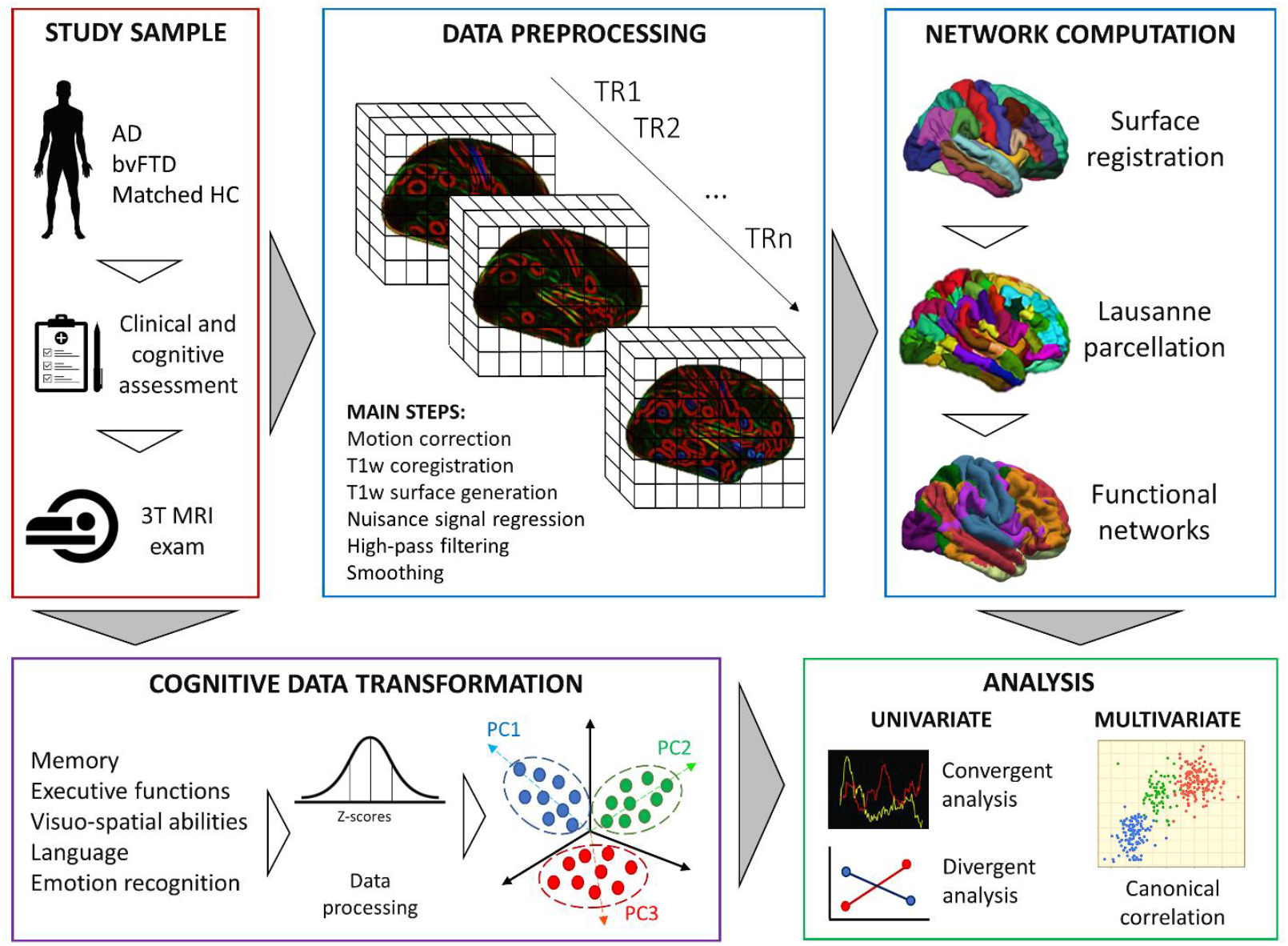
Workflow of the methodology. Patients and controls underwent an extensive cognitive and clinical assessment and a 3T magnetic resonance imaging (MRI) exam. MRI functional data were preprocessed, registered to the subject surface and parcellated according to the Lausanne template. From these parcels, we computed the functional connectivity strength according to the Yeo’s networks template. Each cognitive test score was z-scored and entered in a principal component analysis (PCA) to identify the main cognitive components. Both univariate and multivariate approaches were applied to investigate the relationship between cognition and functional connectivity. Abbreviations: AD: Alzheimer’s disease; bvFTD: behavioral frontotemporal dementia; HC: healthy controls; TR: time repetition. PC: principal component.

### 2.2 MRI acquisition

Rs-fMRI and structural MRI data were acquired on a 3T Philips Achieva system equipped with 8-channel head-coil (University Hospital of Verona, Italy). The following sequences and parameters were used: 2D gradient echo echo-planar imaging (GRE-EPI) sequence for functional connectivity analysis (time repetition; TR/echo time; TE=3000/30ms; flip angle=80°, resolution=3mm isotropic; 48 axial slices; volumes=200); 3D structural T1-weighted (TR/TE=8/3.7ms; flip angle=8°; resolution=1mm isotropic; 180 sagittal slices). Four fMRI volumes with reversed phase encoding directions were acquired for distortion correction purposes. Subjects were instructed to lie still in the scanner, to keep eyes closed but not to fall asleep while images were collected.

### 2.3 Imaging processing and computation

The first 5 scans were removed for stability of the signal. Scans were corrected for distortions using the FMRIB’s Software Library (FSL, fmrib.ox.ac.uk/fsl/) topup tool^26^. Imaging preprocessing was performed according to a previously validated approach used by our group^27^. Specifically, we i) computed motion parameters through a custom preprocessing script; ii) computed the affine registration matrix between rs-fMRI and the T1 image^28^ and iii) registered structural parcellation to the rs-fMRI image through Freesurfer version 6.0 (surfer.nmr.mgh.harvard.edu). The BOLD signal was corrected by regressing out effects of motion (six motion parameters) and mean signal in CSF and left-right white matter (from Freesurfer). The signal was additionally band-pass filtered (0.01 - 0.1 Hz), and scrubbed by removing frames with potential movement artifacts (framewise displacement larger than 0.25 and a DVARS value 1.5×IQR above the third quartile). After this procedure 1 bvFTD patient was excluded due to excessive motion. Additionally, 1 frame preceding each frame with potential movement artifacts was also removed to accommodate temporal smoothing of the signal. Lausanne parcellation^29^ was registered to the native space and used as mask to compute the correlation between the 120 parcels of the template. Finally, for each cognitive network from the Yeo’s template we calculated the average connectivity strength, computed as the correlation average of each parcel included in the network template according to the following formula:

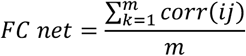

where corr(*ij)* represents the Pearson’s correlation between each pair of parcels belonging to a specific cognitive network from Yeo’s template (DMN, DAN, FPN, and VAN), and m represents the number of all the pairs belonging to the network.

Finally, hippocampal volume was computed through FreeSurfer. For each subject, left and right hippocampal volumes were first corrected for intracranial volume and then averaged to compute a unique hippocampal metric. This metric was used to investigate the association with cognitive components.

### 2.4 Cognitive and clinical assessment

The neuropsychological evaluation included the following tests: Auditory Verbal Learning Test, immediate and delayed recall^30^, Rey–Osterrieth Complex Figure recall^31^, story recall^32^, paired associates learning test (PAL)^33^, digit span backward test^34^, verbal fluency (phonemic and semantic) tasks^35^, Token Test^36^, Trail Making Test part A (TMT-A) and part B (TMT-B)^37,38^, Rey–Osterrieth Complex Figure copy^31^, Reading the Mind in the Eyes^39^, and the 60 Ekman faces tests^40^. Patients’ score at each test was z-transformed based on the performance distribution of the whole sample (patients and age-matched HC). Z-scores for reaction times (i.e., PAL, TMT-A and TMT-B) were inverted for congruency with performance scores of the other tests, i.e., higher scores representing better performance. Due to the high proportion of missing value in the TMT-B, this test was excluded from the PCA analysis, to avoid possible biases. The clinical assessment included the Clinical Dementia Rating (CDR) Scale (global and Sum of Boxes (CDR-SOB) scores)^41^, the Neuropsychiatric Inventory (NPI)^42^, the Frontal Behavior Inventory (only for bvFTD)^43^, and the Instrumental Activities of Daily Living scale (IADL)^44^.

### 2.5 Cognitive components characterization

The subjects × cognitive z-scores matrix was fed into a PCA using the Statistical Package for the Social Sciences (SPSS – Inc., version 23.0. Chicago). We expected the components to be correlated, so an oblique rotation was used, in line with previous literature^22^. Components had to satisfy two criteria: i) the eigenvalues had to be > 1; ii) the percentage of variance accounted for had to be > 5%. We excluded 1 AD and 1 bvFTD patient due to cognitive data missing to avoid possible biases in the PCA computation.

We further compared the cognitive components (referred as cogPC) with cognitive composite scores, computed according to previous published procedure^24^. Z-scores from each test within a specific domain were averaged to compute 5 different composite scores: memory included the Auditory Verbal Learning Test, immediate and delayed recall, the Rey–Osterrieth Complex Figure recall, the story recall, the digit span backward test, and the paired associates learning test; language included the verbal fluency (phonemic and semantic) tasks, and the Token Test ; executive functions included the Trail Making Test part A and part B, visuo-constructional abilities included the Rey– Osterrieth Complex Figure copy and the clock test; emotion recognition included the Reading the Mind in the Eyes, and the 60 Ekman faces tests. We compared each cogPC with the composite scores by means of a linear regression analysis. Different models were computed, each one having cogPC and composite scores as dependent and independent variables, respectively. We further investigated the Spearman’s correlation between each cogPC with clinical outcomes (i.e., IADL, NPI, CDR-SOB and FBI (only in bvFTD) outcomes). Statistical differences among groups in cogPC scores were assessed with the nonparametric Kruskal-Wallis test (AD *vs* bvFTD *vs* HC). Additionally, cogPCs scores were binarized into positive (x>0) and negative (x<0) values. For each group (AD, bvFTD, HC), the number of binarized cases were included in a Chi-squared aimed at investigating differences in the distribution of positive and negative values between groups. Finally, to further characterize cogPCs scores, we investigated the (Spearman’s) relationship between each component and hippocampal volume.

### 2.6 Relationship between cognitive components and cognitive networks

Baseline sociodemographic and cognitive profile of patients and controls were assessed with the Kruskall-Wallis or chi-squared tests as appropriate. A Mann Whitney test was performed to investigate FC network differences between each patient group and HC (AD vs HC; bvFTD vs HC). We investigated both convergent and divergent associations between cogPC scores and network FC in the whole cohort. Statistical analyses and figures were done with Python v.3.

#### 2.6.1 Univariate analysis

Convergence between FC and cognition in the whole cohort was investigated through a robust correlation-wise analysis, aimed at investigating the association between network FC and cognitive components in the whole dataset. Specifically, this analysis was performed between the four cognitive networks from Yeo’s atlas (DMN, FPN, DAN and VAN) with each cogPC scores. Spearman’s correlations between cogPC scores and FC of each network were computed after removing one subject randomly at each step (*leave-and-take-one-out procedure*). In total, we performed 16 steps (corresponding to the exclusion of the 25% of the whole dataset for the last run of the procedure). This procedure was randomly repeated 10000 times to improve the robustness of the result, for a total of 160000 spearman’s correlation values computed for each network-connectivity coupling. The median p-values and Spearman’s coefficients of this convergent correlation-wise analysis distribution were then computed and plotted. To assess the statistical significance of this median r values we implemented a permutation analysis. For each cogPC scores-network coupling we permuted the data 100 times (permutating FC values keeping fixed cogPC scores). At each permutation step we performed 100 random steps excluding until the 25% of the whole dataset, as in the convergent correlation-wise analysis. This procedure led to the same amount of spearman’s (permuted) correlation values (n=160000). Median convergent correlation-wise r values were z-scored according to the mean and standard deviation of this resulting permutation distribution and the corresponding p-values were computed.

Divergent cognitive-connectivity coupling was assessed through a general linear model (GLM), assessing the diagnosis*network interaction for each cogPC. For the interaction analysis, we considered AD, bvFTD and HC as diagnostic factors. For each analysis, we excluded network data points above or below the 1.5 interquartile range. Finally, the same GLM model was repeated only for the patient cohort (AD and bvFTD).

#### 2.6.2 Multivariate analysis

To confirm the relationship between cognitive scores and cognitive networks we applied a canonical correlation analysis (CCA). This approach quantifies the multivariate association between patterns of network connectivity measures and cognitive scores, seeking the maximal correlation between linear combinations of variables in two different sets, i.e., FC and cognitive performance. Cognitive networks showing robust convergent univariate associations (i.e., the convergent correlation-wise analysis) were included as the network dataset. In addition, the visual and sensorimotor networks were included as control networks, as we did not expect a significant association within a low dimensional cognitive space in this cohort, as one would expect for different brain disorder, such as stroke^22,45^. Before CCA, network FC values were z-scored. The five cognitive z-scored composite scores were included as the cognitive dataset (see section 2.5). CCA modes exhibiting a significant correlation between variates from the whole group were compared between groups through analysis of variance (ANOVA), testing both main effects and interactions.

## 3. Results

Twenty-two AD and 23 bvFTD patients were included in the study. A sample of 20 age-matched individuals was included as control group. Patients and HC were comparable for age (p=0.904), education (p=0.910), and gender (p=0.344). As expected, the MMSE scores was significantly lower in patients compared to HC (p<0.001), without differences between AD and bvFTD (p>0.05). Compared to HC, patients showed lower cognitive scores in all the cognitive tests (p<0.002 for all scores). As expected, bvFTD exhibited lower performance compared to AD for the phonemic fluency test, the reading the mind in the eyes and the 60 Ekman faces tests (**Table 1**). bvFTD compared to AD showed more severe behavioral disturbances (NPI: p<0.001; FBI: p=0.001) and disease severity (IADL: p=0.012; CDR-SOB: p=0.04). Both bvFTD and AD showed significantly lower hippocampal volumes compared to HC (around 20% of volume reduction for both patient groups; p<0.001).

**Table 1.** Baseline sociodemographic and cognitive profile of patients and controls.

|  | **HC**<br>**n=20** | **AD**<br>**n=22** | **bvFTD**<br>**n=23** | **p** |
| --- | --- | --- | --- | --- |
| Age | 72 ± 6 | 72 ± 6 | 71 ± 10 | .904 |
| Sex (% Female) | 50% | 55% | 57% | .910 |
| Education | 11 ± 5 | 9 ± 4 | 9 ± 4 | .105 |
| MMSE | 29 ± 2 | 21 ± 2 * | 22 ± 4 * | <.001 |
| Left hippocampus | 3724 ± 339 | 2892 ± 433 * | 2916 ± 684 * | <.001 |
| Right hippocampus | 3794 ± 326 | 2951 ± 451 * | 3079 ± 708 * | <.001 |
| **Cognitive tests** |  |  |  |  |
| RAVLT – immediate | 46 ± 7 | 20 ± 7* | 21 ± 7* | < .001 |
| RAVLT – recall | 10 ± 2 | 1.05 ± 1.50* | 2.43 ± 2.63* | < .001 |
| Episodic memory | 13 ± 3 | 2.43 ± 2.18* <sup>a</sup> | 3.89 ± 3.80* | < .001 |
| PAL | 34 ± 16 | 166 ± 28* | 146 ± 54* <sup>a</sup> | < .001 |
| ROCF - recall | 15 ± 4 | 2.31 ± 3.15* <sup>b</sup> | 5.17 ± 4.57* | < .001 |
| Backward digit span | 4 ± 1 | 2.59 ± 1.87 | 1.78 ± 1.98* | 0.002 |
| Clock test | 12 ± 1 | 5.59 ± 3.39* | 7.35 ± 3.31* | < .001 |
| ROCF - copy | 30 ± 4 | 22 ± 10* <sup>b</sup> | 20 ± 9* | < .001 |
| TMT-A | 46 ± 10 | 120 ± 85* <sup>b</sup> | 134 ± 94* | < .001 |
| TMT-B | 129 ± 46 | 387 ± 194* <sup>c</sup> | 267 ± 127* <sup>d</sup> | < .001 |
| Phonemic fluency test | 35 ± 8 | 24 ± 10* | 13 ± 8 *# | < .001 |
| Semantic fluency test | 40 ± 6 | 19 ± 8* | 16 ± 8* | < .001 |
| Token test | 34 ± 2 | 28 ± 5* | 25 ± 6* | < .001 |
| RMET | 20 ± 3 | 16 ± 4* | 12 ± 4*# | < .001 |
| EK-60F | 47 ± 5 | 39 ± 8* | 27 ± 10*# | < .001 |

\* variables statistically different from HC; # variables significantly different between bvFTD and AD.
a) data from 22 subjects; b) data from 21 subjects; c) data from 11 subjects; d) data from 8 subjects.
Abbreviations: MMSE: Mini Mental State Examination; RAVLT: Rey Auditory Verbal Learning test; PAL: paired associates learning test; ROCF: Rey–Osterrieth Complex Figure; TMT: Trail Making Test; RMET: Reading the mind in the eyes test; EK-60F: Ekman 60 Faces Test

### 3.1 Cognitive components

The PCA analysis revealed 3 main cognitive components in the whole dataset. A first component (cogPC1) accounted for around 63% of the variance across all tests and subjects. A second factor (cogPC2) accounted for around 11% of variance, while a third factor (cogPC3) accounted for more than 7% of variance (**Figure 2**). In the whole dataset, the first component was significantly associated with hippocampal volume (rho=.649; p<0.001), while the second and third components were not (p>0.080).

**Figure 2.**
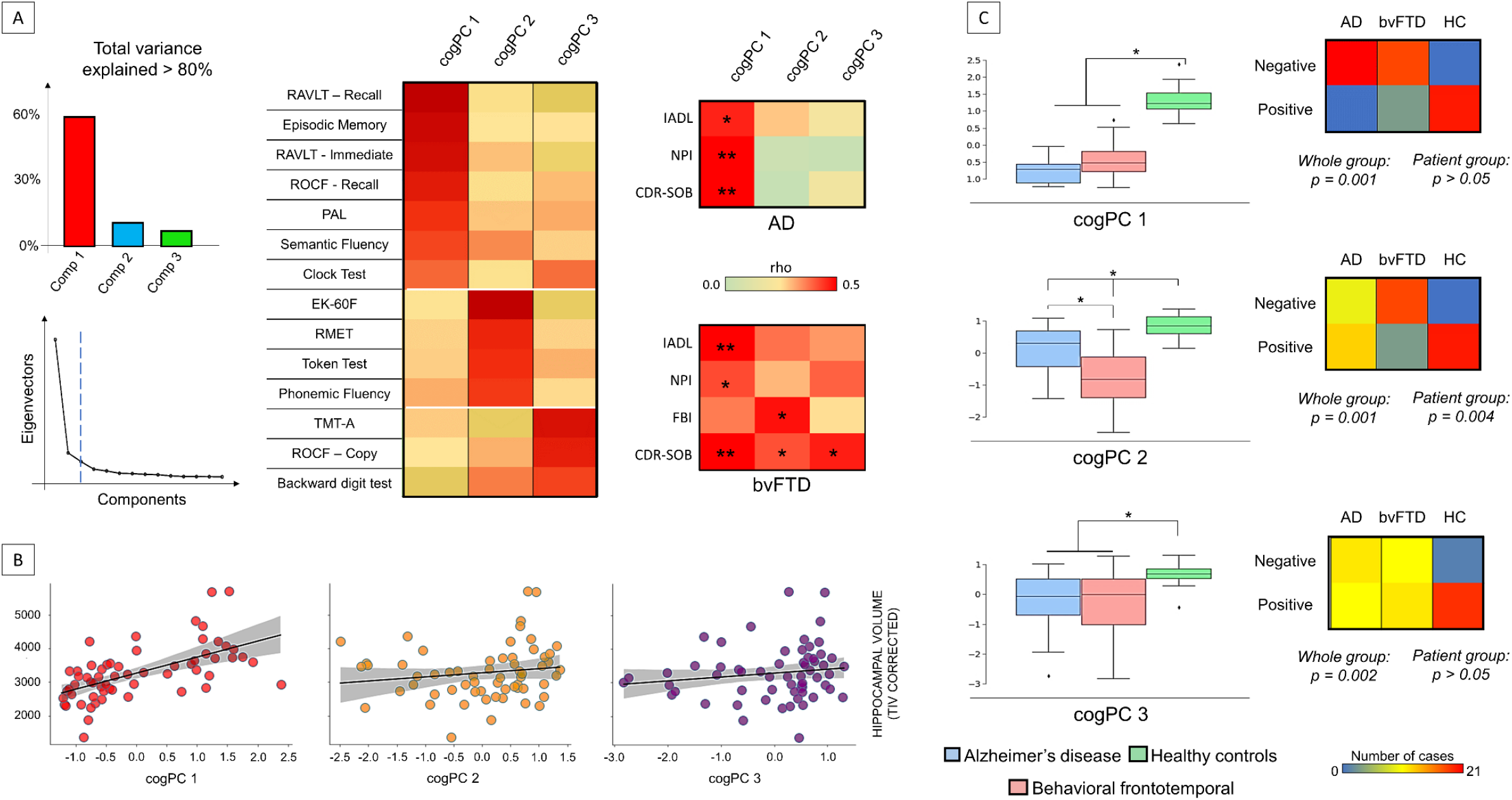
Principal component analysis of cognitive scores. Panel A: cognitive z-scores were entered in a principal component analysis (PCA), revealing 3 main components (cogPC) explaining more than 80% of variance (left panel). The matrices in the middle panel report the loadings of the PCA. The right matrices report the correlations between cognitive component scores and clinical outcomes in each patient group separately, showing specific-disease involvement. Panel B: the first component was related with hippocampal volume in the whole sample (p<0.001), while not significant relationships were reported between hippocampus and the other components scores (p>0.05). Abbreviations: AD: Alzheimer’s Disease; bvFTD: behavioral variant frontotemporal dementia; ROCF: Rey–Osterrieth complex figure test; RAVLT: Rey Auditory Verbal Learning Test; PAL: paired associative test; EK-60F: The Ekman 60-Faces; RMET: Reading the Mind in the Eyes; NPI: neuropsychiatric inventory; FBI: frontal behavioral inventory; IADL: Instrument activity daily life; CDR-SOB: clinical dementia rating – sum of boxes; TIV: total intracranial volume. Panel C: significant differences were reported in patients compared to controls in all components. BvFTD patients showed lower scores compared to AD in the second cognitive component (left side). Differences in binarized component scores were reported among groups for all the components (right side). When considering only patient groups, a significant difference was reported for the second component.

The results for the linear regression analysis comparing each cogPC with the composite z-scores are reported in **Table 2**. In AD and bvFTD, the first component, showed a significant association with the memory composite (p<0.001 for both), while the remaining composite scores were not significant (p>0.100 and p>0.20, respectively). The second component in AD showed a strong association with language and emotion recognition composite scores (both p<0.001). This result was echoed in bvFTD (emotion and language p<0.001), with an additional effect for memory (p=0.02). Finally, in AD the third component showed a significant association with visuo-spatial abilities (p<0.001) and language composite scores (p=.012). In bvFTD the third component showed a strong association with visuo-spatial abilities and executive composite scores (p<0.001). Based on these results, we interpreted the cogPCs as memory component (cogPC1), emotion-language component (cogPC2) and visuo-spatial attentional component (cogPC3) respectively.

**Table 2.** Linear regression analysis for composite and component scores from the principal component analysis.

| Composite scores | Beta | p-value | Beta | p-value |
| --- | --- | --- | --- | --- |
|  | AD |  | bvFTD |  |
| **cogPC 1** | R<sup>2</sup>=0.644, p=.05 |  | R<sup>2</sup>=0.873, p<.001 |  |
| Memory | *0.955* | *< 0.001* | *1.134* | *< 0.001* |
| Executive functions | -0.056 | 0.783 | -0.137 | 0.254 |
| Visuo-spatial abilities | 0.219 | 0.344 | 0.026 | 0.852 |
| Language | -0.091 | 0.683 | -0.184 | 0.354 |
| Emotion recognition | -0.382 | 0.107 | -0.145 | 0.431 |
| **cogPC 2** | R<sup>2</sup>=0.964, p<.001 |  | R<sup>2</sup>=0.975, p<.001 |  |
| Memory | -0.111 | 0.095 | -0.142 | *0.019* |
| Executive functions | -0.039 | 0.547 | -0.034 | 0.525 |
| Visuo-spatial abilities | -0.108 | 0.150 | -0.061 | 0.337 |
| Language | *0.525* | *< 0.001* | *0.596* | *< 0.001* |
| Emotion recognition | *0.738* | *< 0.001* | *0.557* | *< 0.001* |
| **cogPC 3** | R<sup>2</sup>=0.893, p<.001 |  | R<sup>2</sup>=0.904, p<.001 |  |
| Memory | 0.044 | 0.688 | -0.064 | 0.555 |
| Executive functions | 0.140 | 0.216 | *0.412* | *0.001* |
| Visuo-spatial abilities | *0.696* | *< 0.001* | *0.691* | *< 0.001* |
| Language | 0.340 | *0.012* | 0.049 | 0.733 |
| Emotion recognition | -0.169 | 0.130 | -0.120 | 0.452 |

All these cognitive components were different among groups. Post-hoc analysis showed that cogPC1 and cogPC3 were different in both patient groups compared to HC (p<0.002). By contrast, we reported a gradient in the cogPC2, with HC showing the greatest score, followed by AD and then by bvFTD (p=0.031). The difference between AD and bvFTD was significant (p=0.015) (**Figure 2**). The same between-group differences were observed for the cogPCs scores binarized into positive and negative values (**Figure 2**).

When investigating the relationship between cogPCs with clinical outcomes, the first component showed a strong relationship with IADL, NPI and CDR-SOB in both patient groups. Lower cognitive scores were positively associated with IADL (indicating greater functional disability) and negatively with NPI and CDR-SOB (indicating greater behavioral disturbances and higher disease severity). Notably, in AD the other two components were unrelated with clinical outcomes, while in bvFTD we found an association between the second component with both behavioral disturbances (FBI) and disease severity (CDR-SOB), and the latter with disease severity (CDR-SOB). These associations were positive, indicating a relation between lower scores and greater disabilities (**Figure 2**).

### 3.2 Brain functional networks and connectivity-cognitive coupling

As shown in the **Supplementary Figure S1**, VAN connectivity was lower in bvFTD compared to HC (p=0.01), while connectivity values for DAN, FPN and DMN were lower in AD compared to HC (p<0.04).

The convergent correlation-wise analysis revealed a robust and stable significant correlation between the first two cognitive components with the DMN and VAN (**Figure 3**). Specifically, we reported a median rho correlation of 0.303 between the memory component and the DMN (pmedian DMN=0.026), indicating that lower cognitive performance was associated with lower FC. Similarly, the emotion-language component showed a median rho correlation around 0.30 for both the DMN and VAN (pmedian DMN=0.023; pmedian VAN=0.043). These median correlation values compared to a permutation distribution showed statistical significance for the DMN (memory component: p=0.016; emotion-language component p=0.042) and near to the significance level for the VAN (emotion-language component p=0.056) (**Figure 3**). By contrast, DAN and FPN showed no significant association with any component. Similarly, the visuo-spatial attentional component showed no significant correlation with the cognitive networks (**Figure 3**).

**Figure 3.**
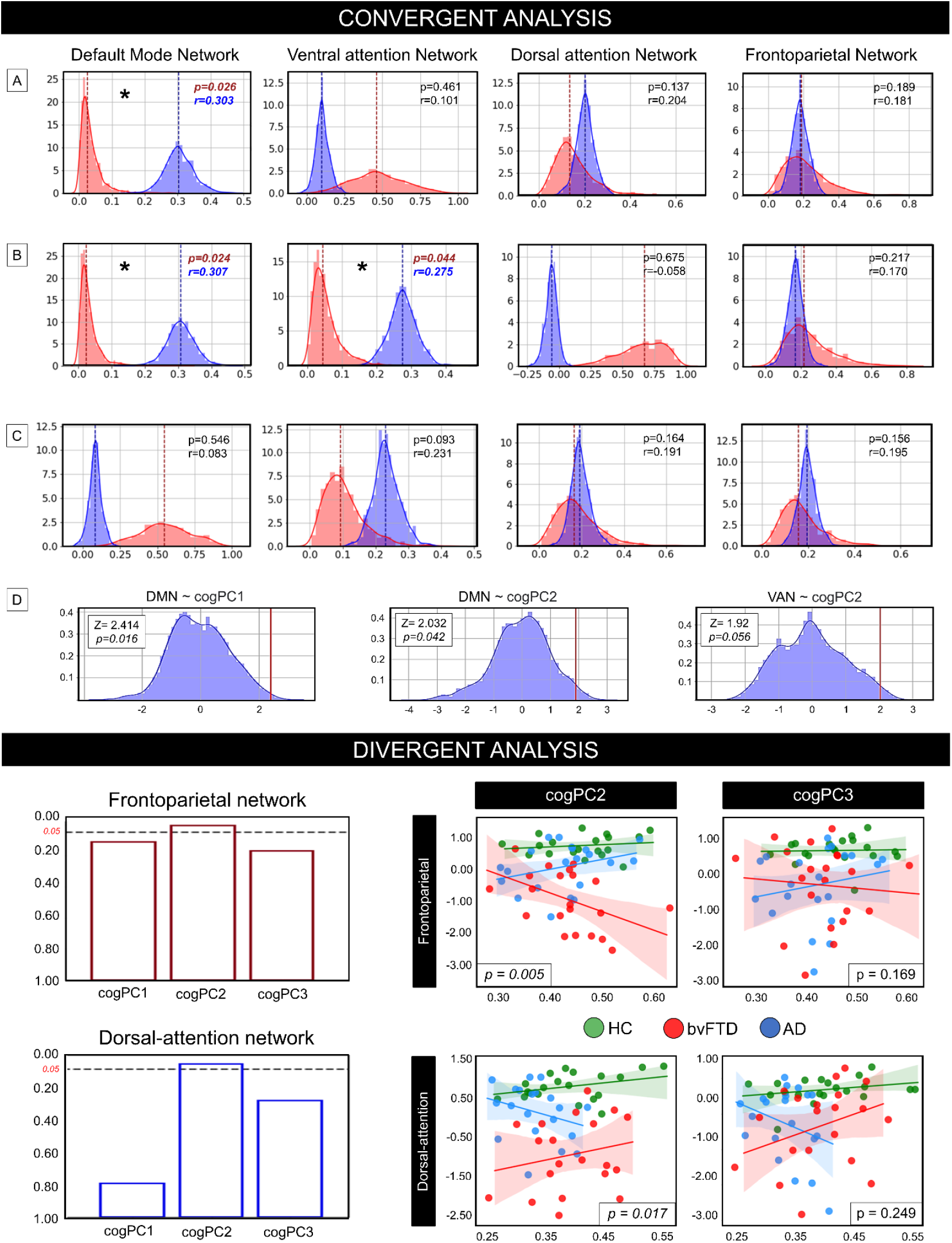
Univariate convergent correlation-wise and divergent analyses between cognition and connectivity. Top panel: Spearman’s correlation and p-values for the convergent correlation-wise analysis in the whole dataset. Correlations were computed for each cognitive network (default mode network, ventral and dorsal attention network, and frontoparietal network) and cognitive component scores. Panel A: correlation values distribution between connectivity and cogPC1 (memory component); Panel B: correlation values distribution between connectivity and the cogPC2 (emotion-language component). Panel C: correlation values distribution between connectivity and the cogPC3 (visuo-spatial attentional component). Panel D: permutation analysis for the significant convergent median p-values are reported (bottom panel). Bottom panel: Interaction effect analysis between network, diagnosis and cognition for each cognitive component and functional network. A significant divergent effect between diagnostic group and network was reported for the non-memory cognitive components with the attentional networks. Top panel: bar plots of the diagnosis*cognitive component significance interaction with the attentional networks. Bottom panel: scatter plot of the interaction for the non-memory components and the attentional networks. Analyses were performed after exclusion of network data outliers.

The interaction analysis showed that the lack of a significant relationship between cognition and the DAN/FPN in the whole dataset was due to a divergent coupling effect between groups. Specifically, after the exclusion of two outlier data points the GLM showed a significant diagnosis*cogPC2 for the DAN (z=-2.389; p=0.029) but not for the interaction between diagnosis and cogPC3 (z=-1.152; p=0.249) (**Figure 3**). In bvFTD, lower cognitive scores were associated with lower FC, while the opposite was seen in AD. Similarly, for the FPN, after the exclusion of one outlier data point, we reported a significant diagnosis*cogPC2 effect (z=-2.809; p=0.005), but the relationship was reversed: lower scores were linked with lower FC in AD and the opposite in bvFTD. The cogPC3 showed no significant interaction (z=-1.376; p=0.169) (**Figure 3**). No interactions were reported between diagnosis and cogPC1 with both DAN and FPN (p>0.10 for all the analysis). DMN and VAN showed no divergent patterns with cogPC scores (p>0.30), in line with the robust convergent association of these networks. These results were confirmed when considering only patients (AD and bvFTD; **Supplementary Figures S2)**.

Within-diagnosis network-cognitive interaction effects, that is FPN*DAN interactions separately for AD and bvFTD, confirmed the previous results. We found a divergent within-diagnosis effect between cogPC2 with FPN and DAN in both cohorts (DAN*FPN interaction effect p=0.012 for both patient groups). A trend was reported for the DAN*FPN interaction with cogPC3 (AD: p=0.078; bvFTD: p=0.128) (**Supplementary Figures S3**).

### 3.2 Multivariate association between cognitive performance and network connectivity

We considered cognitive networks showing a robust univariate association with cognition (i.e., DMN and VAN), as the CCA seeks the maximal correlation between linear combinations of variables in two different sets. We identified 2 pairs of modes that significantly correlated the network variables and cognitive performance (mode1: r=0.51, p<0.001; mode2: r=0.48; p<0.001). The first mode mainly loaded on memory, language, and emotion recognition, while the corresponding network mode loaded on the DMN. The second mode mainly loaded on language and emotion recognition on the cognitive side, and on VAN on the corresponding network side (**Figure 4**). ANOVA showed a significant difference for both cognitive modes (p<0.001) (**Figure 4**). Post-hoc Tukey’s test revealed that the first mode was significantly different between HC and both disease groups (AD: p<0.001; bvFTD: p<0.001), while AD and bvFTD showed no significant differences. The second mode were different between groups (p<0.001), mirroring cogPC2 result in bvFTD, as this group showed significant differences compared to both AD (p=0.033) and HC (p<0.001). No differences were reported between AD and HC (p=0.285).

**Figure 4.**
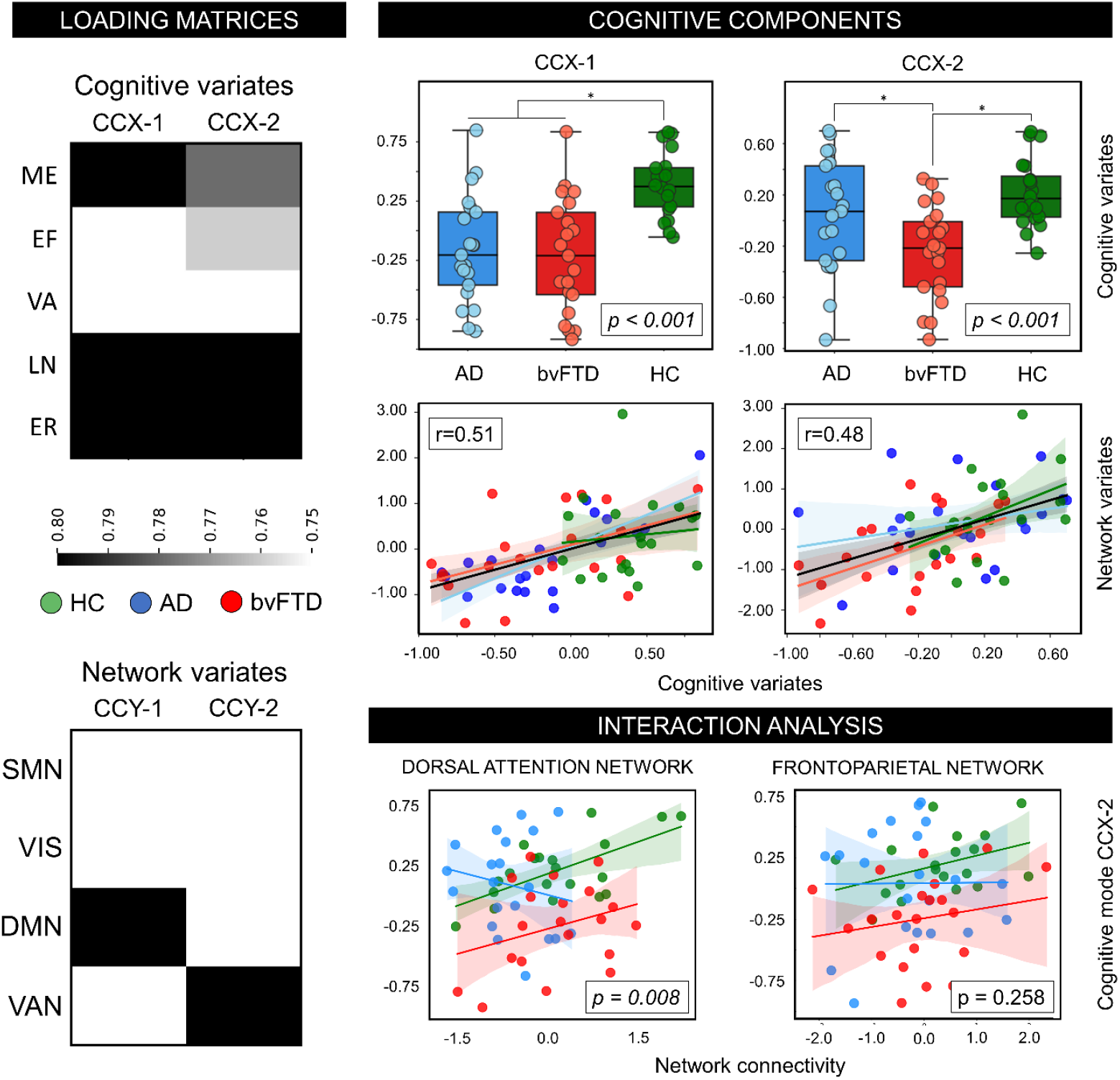
Multivariate canonical correlation analysis. Two pairs of mode showed the maximal correlation between cognitive composite scores and network connectivity. The first pair of mode loaded on memory, language and emotion recognition composite scores (cognitive dataset), and DMN (network dataset); the second pair of mode loaded mainly on the language and emotion scores and the VAN (left panel). Values and loadings from the first mode were inverted to improve the comparison with the second mode. Values associated with the cognitive modes were different among groups (right panel). A divergent group effect was reported between these cognitive modes and DAN connectivity (panel bottom-right). Abbreviations: DMN: default mode network; EF; executive functions; ER: emotion recognition; LN: language; ME: memory; SMN: sensorimotor network; VA: visuo-spatial abilities; VAN: ventral network; VIS: visual network.

Finally, we tested the GLM interaction model between the second mode (echoing the cogPC2), between groups with DAN and FPN connectivity, after removing outliers according to the 1.5 interquartile range. We confirmed a divergent association between this cognitive mode with DAN (p=0.008), while FPN did not show a significant effect (p=0.258) (**Figure 4**).

## 4. Discussion

The core results of this study are: (a) a low dimensional cognitive space in the aging and age-related pathology continuum; (b) both divergent and convergent FC patterns linked with this low cognitive space. These findings could shed light into the relationship between cognitive alterations and brain alterations in AD and bvFTD, suggesting possible detrimental and compensatory mechanisms.

### 4.1 Low cognitive dimensional space

In this study we identified three main cognitive components across normal aging, AD and bvFTD explaining more than 80% of variance. The first memory component explained the largest amount of variance in our sample and was significantly related to hippocampal volumes, congruently with previous literature^46^. This component was also associated with the clinical severity in both dementia groups and may therefore represent a common neuropsychological impairment across AD and bvFTD^47,48^. The second component was represented by a language-emotion factor that was involved in both AD and bvFTD but was more impaired in the latter group and linked with both clinical severity and behavioral disturbances only in this cohort. This component may therefore capture a neuropsychological feature specific to bvFTD. Emotion and language are indeed two functions highly impacted in this disorder^49,50^ and their inclusion within a common factor is not surprising as language plays a fundamental role in emotion. Previous researches highlighted that access to the meaning of emotional words (i.e., with emotional meaning) is a fundamental component to understand emotional facial expressions^51^. Finally, we identified a third component, mainly loading to visuo-spatial executive tests. This component, although altered in both patient groups, was linked with the clinical severity only in bvFTD, again suggesting that this factor may capture a bvFTD-specific neuropsychological feature. Overall, these findings suggest a low-dimensional cognitive pattern within the aging and age-related pathology continuum, where only three cognitive dimensions explained a large amount of variance of cognitive outcomes. Previous research identified a similar low dimensional pattern in stroke; despite the great heterogeneity of brain lesions, it has been suggested a low dimensional pattern of cognitive deficits, involving three main different components^22^. These factors might help to identify a low space for behavioral phenotypes in neurodegeneration, moving beyond the classical “composite” score approach. Indeed, the latter approach is critically dependent upon the *apriori* definition of the cognitive domains measured by the different tests, thereby neglecting the frequent co-occurrence of deficits within and across domains.

### 4.2 Divergent and convergent relationships between cognition and connectivity

This low dimensional cognitive manifold was linked with the FC pattern of higher-order cognitive networks. FC of the DMN was positively associated with memory and emotion-language scores in the whole cohort, with no significant interaction between groups. The association between DMN and memory is in line with a vast body of literature^52^, while the association between DMN and emotion-language suggest that this network plays an important role for constructing discrete emotional experiences^53^. Additionally, we reported a robust association between the VAN and the emotion-language scores, with no significant interaction between groups, linking the VAN with social functioning^54^. Previous studies highlighted that the VAN can be briefly activated by external stimuli of behavioral relevance^19^, suggesting that stimuli encoded in VAN areas are defined also by emotional experience^55^. These results were confirmed by the multivariate analysis, showing a robust linear relationship between DMN with a cognitive mode represented by memory, language, and emotion recognition, and between a cognitive mode mirroring the cogPC2 with VAN connectivity (as shown in **Figure 4**). Although DMN and VAN showed a selective vulnerability in AD and bvFTD, respectively, these associations suggest shared multi-dimensional network mechanisms between these disorders, congruent with the network-cognitive relationships observed in physiological conditions. The link between DMN with memory and VAN with emotion-language might be ubiquitous within the aging-pathology continuum, although bvFTD showed higher levels of deficit in both language-emotions and VAN connectivity, suggesting that failure in these networks is associated with the decline of memory and social domains.

Along with these commonalities, we reported divergent connectivity-cognitive couplings, confirmed by both the univariate and the multivariate analysis. For both DAN and FPN we reported a divergent pattern between the emotion-language component and diagnosis. This effect was confirmed by the analysis performed in the patient cohort, in addition to a significant divergent effect between the DAN and the visuo-spatial component. Overall, this pattern suggests that divergent network-level effects might emerge as consequence of aberrant connectivity observed in the primary affected networks in these disorders. In the last years, it has been well established the role of DAN as a “network gate” facilitating top-down attention processing by suppressing VAN signals to exclude irrelevant bottom-up information^18,56,57^. We speculate that in bvFTD, given normal connectivity of the DAN but reduced connectivity of the VAN, the positive relationship between DAN and non-memory cognitive components may reflect compensatory neural mechanisms for attentional/emotional/language processing typifying this disorder. Indeed, DAN and VAN dynamically interact to control the information to be processed^58,59^ indicating that DAN connectivity in bvFTD might compensate VAN failure. On the other side, AD showed the opposite pattern, i.e., reduced connectivity of the DAN and normal VAN pattern. This may imply in AD a defective role of DAN in facilitating top-down processes inhibiting irrelevant information as well as reduced regulation of attentional networks (i.e., VAN and FPN). Thus, a less flexible dynamic response might result in a less efficient maintenance of cognitive set. A similar divergent pattern between connectivity, cognition and diagnosis was observed for the FPN and the language/emotion cognitive component. In this case the association was reversed. FPN was significantly reduced in AD compared to HC, suggesting that this pattern might highlight residual functionality, that is patients with lower FC have worse cognition, while those with higher cognition show relatively preserved cognitive performance. A coupled activity between DMN and FPN supports cognitive demand for goal-directed task^60^. Moreover, the positive association between these two networks was associated to between-network compensatory mechanisms in mild cognitive impairment patients^61^, indicating that FPN connectivity might sustain DMN failure. Again, in bvFTD this pattern was reversed, suggesting an emerging defective role of FPN over cognitive functions. Overall, these results point to the presence of both common and divergent patterns among AD and bvFTD patients, suggesting that cognitive alterations are distributed among a connectivity dysfunctional gradient, which may reflect reduced variability in network dynamics. FC of distant cognitive networks represents a dynamic process and might influence cognitive demands and neural resources, which may be reflect either compensation or network failure.

### 4.3 Limitations and strength

This study has several limitations. First, our sample size was relatively small, although clinical and demographical characteristics of patients and controls were well matched. Second, we did not collect amyloid and tau biomarkers, thus we could not investigate whether FC-cognitive associations were driven by molecular pathology, as suggested by the cascading network failure hypothesis^62^. According to this model, tau-associated local network failure may be followed by a global compensatory phenomenon associated with Aβ^62^. Future studies examining the relationship between molecular pathology and divergent networks connectivity will allow for a more nuanced clarification of the relationship between cognition and FC.

Besides these limitations, this study has two main strengths: (1) investigating for the first time the cognitive dimensional space within the aging and age-related pathology continuum through a dimensionality reduction approach, and (2) assessing the univariate and multivariate relationships between this new cognitive space and neural networks connectivity. Future studies could further confirm these patterns. The identification of divergent and shared neural mechanisms across neurodegenerative diseases would increase our understanding of network dynamics. Moreover, these findings would be useful to optimize non-invasive electric brain network stimulation intervention,^7^ improving target selection for stimulation protocols aiming at rehabilitating specific cognitive functions.

## 5. Conclusions

In conclusion, a PCA approach revealed a low cognitive dimensional space across aging, AD and bvFTD. Cognitive deficits in patients are more accurately described by correlated deficit components rather than the collection of individual scores. We identified a few components that were consistent across different cohorts. The associated network coupling of the identified components showed both convergent and divergent patterns, suggesting both possible detrimental and compensatory effects, which might help to drive new effective interventions^7^.

## Acknowledgments

This work was supported by the Italian Ministry of Health (Giovani Ricercatori grant GR2011– 02349787, Ricerca Corrente).

## Data statement

The datasets generated and/or analyzed during the current study are available from the corresponding author on reasonable request. The dataset is publicly available at the following DOI: 10.17632/s3t9mptvh8.1.

## Ethics approval and consent to participate

The study was approved by the local ethics committee of the IRCCS Istituto Centro San Giovanni di Dio Fatebenefratelli in Brescia (Italy) (Number 43/2014). Written informed consent was obtained from all participants. ClinicalTrials.gov identifier NCT03422250.

## List of abbreviations

AD: Alzheimer’s disease
CCA: canonical correlation analysis
BOLD: blood oxygen level dependent
bvFTD: behavioral variant frontotemporal dementia
DAN: dorsal attentional network
DMN: default mode network
FC: functional connectivity
FPN: frontoparietal network
PCA: principal component analysis
Rs-fMRI: resting-state functional magnetic resonance imaging
VAN: ventral-attentional network

---

**Figure Supplementary S1.**
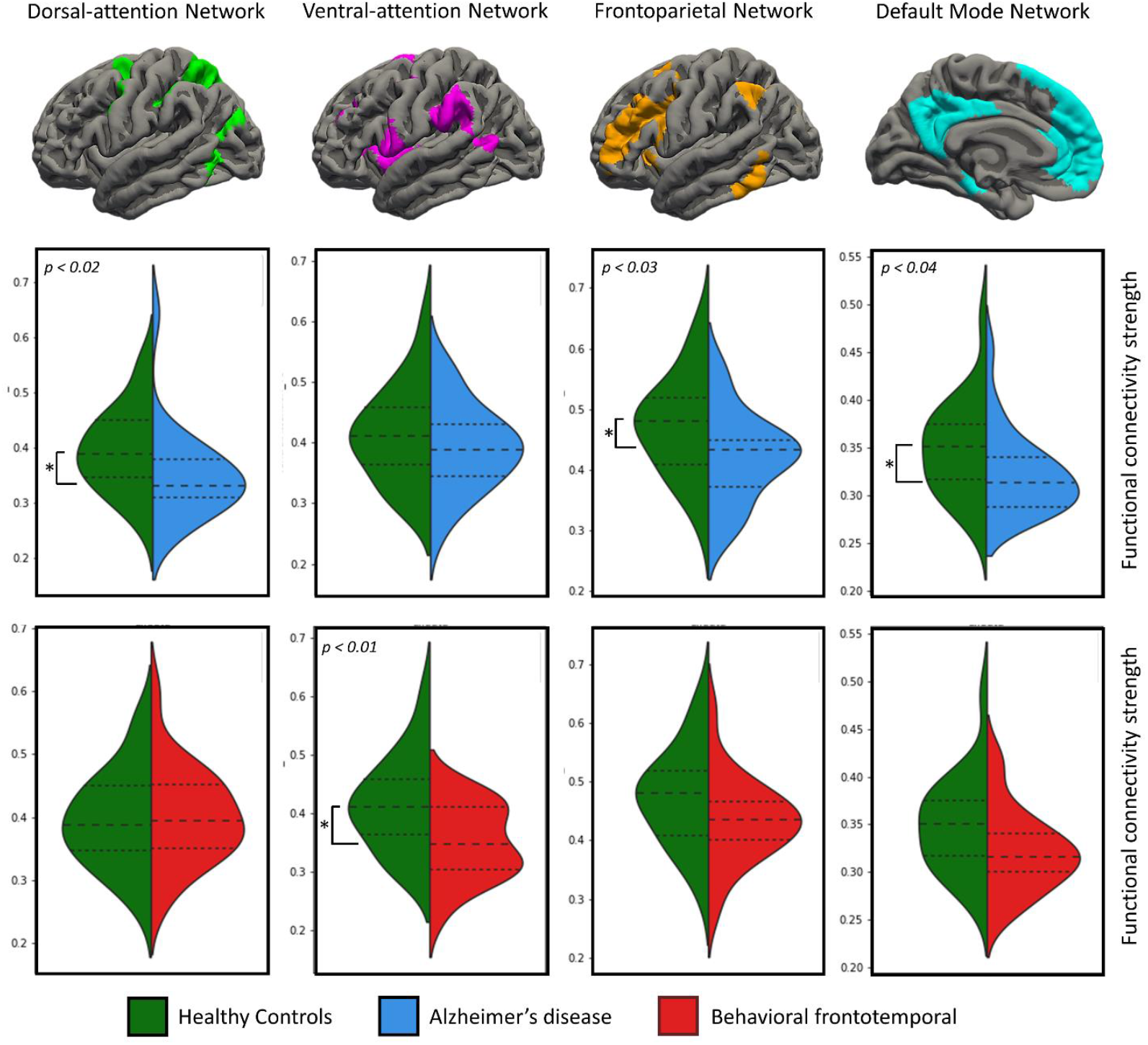
Network differences between patients and controls. Different involvement in AD and bvFTD was reported compared to controls. AD showed lower connectivity in the dorsal-attention network, frontoparietal network and default mode network (top panel). Compared to controls, bvFTD showed reduced connectivity only in the ventral-attention network (bottom panel).

**Supplementary Figure S2.**
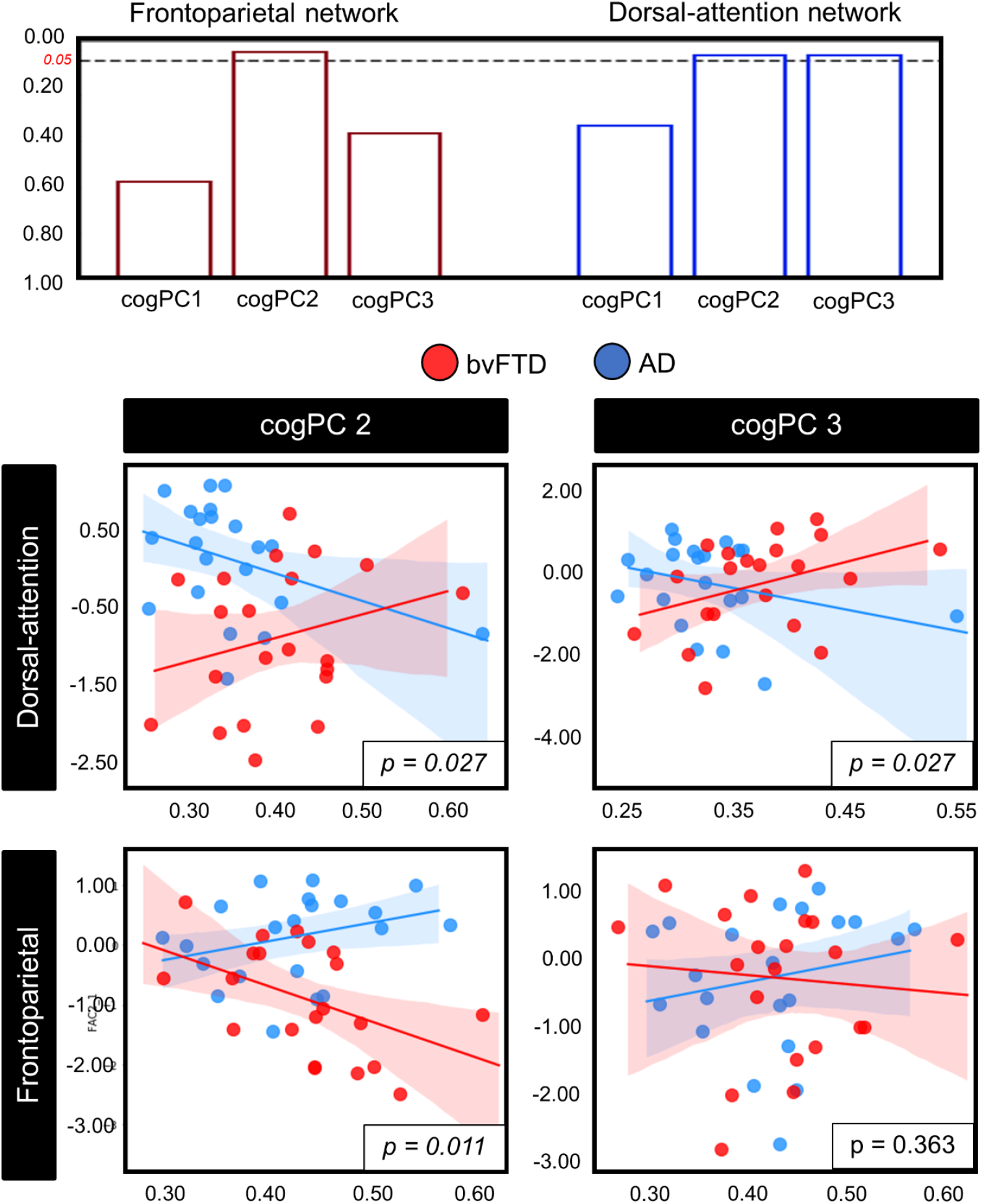
Relationship between connectivity and cognitive components. The main Interaction effect analysis between network, diagnosis and cognition was replicated in the dementia cohort, excluding healthy controls. Top panel: bar plots of the diagnosis*cognitive component significance interaction with the attentional networks. Bottom panel: scatter plot of the interaction for the non-memory components and the attentional networks.

**Figure Supplementary S3.**
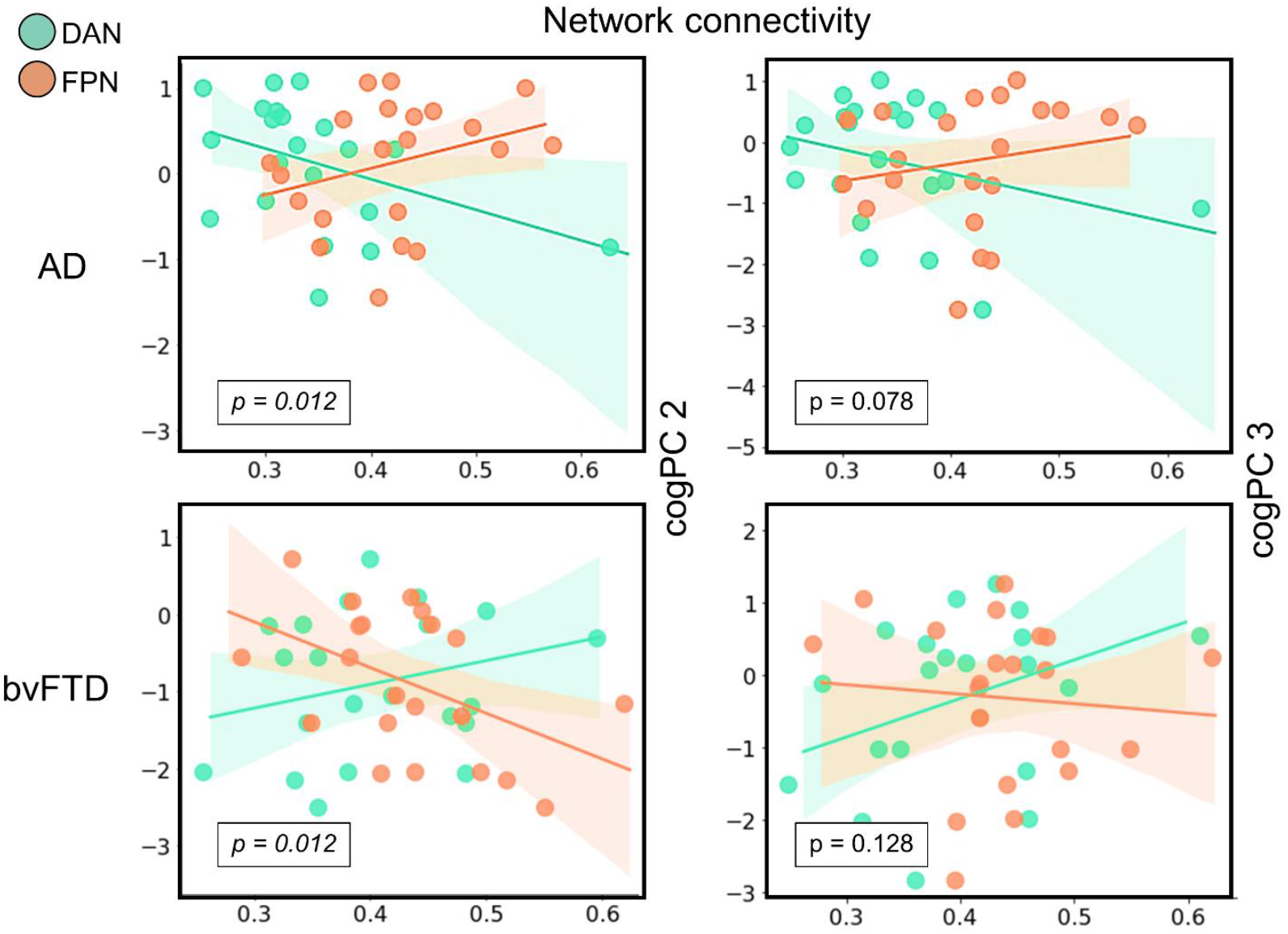
Within-diagnosis divergent cognitive-network coupling. A within-diagnosis cognitive * networks (frontoparietal – FPN – and dorsal attention – DAN) analysis showed similar divergent network-cognitive coupling patterns in both Alzheimer’s disease (AD) and behavioral frontotemporal (bvFTD) patients. Significant divergent effects were reported between the emotion-language component (component 2) and the attentional networks.

